# Interactions of SARS-CoV-2 protein E with cell junctions and polarity PDZ-containing proteins

**DOI:** 10.1101/2021.12.04.471219

**Authors:** Yanlei Zhu, Flavio Alvarez, Nicolas Wolff, Ariel Mechaly, Sébastien Brûlé, Benoit Neitthoffer, Sandrine Etienne-Manneville, Ahmed Haouz, Batiste Boëda, Célia Caillet-Saguy

## Abstract

The C-terminus of the severe acute respiratory syndrome coronavirus 2 (SARS-CoV-2) protein E contains a PBM (PDZ binding motif) targeting PDZ (PSD-95/Dlg/ZO-1) domains identical to the PBM of SARS-CoV. The latter is involved in the pathogenicity of the virus. Recently, we identified ten human PDZ-containing proteins showing significant interactions with SARS-CoV-2 protein E PBM. We selected several of them involved in cellular junctions and cell polarity (TJP1, PARD3, MLLT4, LNX2) and MPP5/Pals1 previously shown to interact with SARS-CoV E PBM. Targeting cellular junctions and polarity components is a common strategy by viruses to hijack cell machinery to their advantage. In this study, we showed that these host PDZ domains TJP1, PARD3, MLLT4, LNX2 and MPP5/PALS1 interact in a PBM-dependent manner *in vitro* and colocalize with the full-length E protein *in cellulo*, sequestrating the PDZ domains to the Golgi compartment. We solved three crystal structures of complexes between human LNX2, MLLT4 and MPP5 PDZs and SARS-CoV-2 E PBM highlighting its binding preferences for several cellular targets. Finally, we showed different affinities for the PDZ domains with the original SARS-CoV-2 C-terminal sequence containing the PBM and the one of the beta variant that contains a mutation close to the PBM. The acquired mutations in E protein localized near the PBM might have important effects both on the structure and the ion-channel activity of the E protein and on the host machinery targeted by the variants during the infection.

## 1 Introduction

Currently, there have been more than 263 million individuals infected by SARS-CoV-2, including more than 5.2 million deaths (https://covid19.who.int/). While significant constant advances are made in understanding this virus, the knowledge of the SARS-CoV-2 proteins interactions with host cell proteins is still limited.

Targeting multiple cellular PDZ (PSD-95/Dlg/ZO-1)-containing proteins through short linear PDZ-binding motifs (PBMs) is a common strategy used by viruses to facilitate their viral replication and dissemination to new hosts (Javier and Rice, 2011). Indeed, PDZ proteins are involved in processes of particular interest in viral infection: cell junctions formation, cell polarity establishment and immune system signaling (Javier and Rice, 2011; James and Roberts, 2016; Gutiérrez-González and Santos-Mendoza, 2019). The two SARS-CoV-2 viroporins, protein E and 3a contain a C-terminal PBM targeting specific PDZ domain-containing proteins (Castaño-Rodriguez et al., 2018; Caillet-Saguy et al., 2021).

PDZ domains are a large family of protein–protein interaction domains widespread in the human proteome (Luck et al., 2012). PBMs are mainly located at the C-terminus of target proteins and interact directly with PDZ domains. They are classified into three types: type I PBM (-X-S/T-X-ϕ_COOH_), type II PBM (-X-ϕ-X-ϕ_COOH_) and type III PBM (-X-D/E-X-ϕ_COOH_), with ϕ signifying a hydrophobic residue. The C-terminal PBM sequence of SARS-CoV E protein is of type II (-DLLV_COOH_) and has been identified as a virulence factor (Jimenez-Guardeño et al., 2014). It is likely that the abilities to target PDZ proteins also make significant contributions to the pathogenesis of the E protein of SARS-CoV-2. Indeed, the E protein is highly conserved with 94.7% identity between SARS-CoV and SARS-CoV-2 and their PBMs are strictly conserved.

Protein E is a small transmembrane protein of 75 residues involved in several phases of the virus life cycle, such as assembly, budding, envelope formation, and pathogenesis (Schoeman and Fielding, 2019). Protein E has a hydrophobic helical transmembrane domain of 30 residues flanked by an N-terminal domain of 8 residues and a C-terminal domain of 37 residues (Fig. 1A) (Mandala et al., 2020). It is present in virions in small quantities whereas it is very abundant in infected cells at the level of the Intermediate Compartment between the Golgi and the Endoplasmic Reticulum (ERGIC) where it actively participates in the budding, morphogenesis and trafficking of the virus. A cytoplasmic orientation of the C-terminal domain and a luminal orientation for the N-terminal domain has been reported (Nieto-Torres et al., 2011; Duart et al., 2020). The E protein can oligomerize via its transmembrane domain into a homopentameric ion channel called viroporin that inserts into the host cell endomembrane system (Li et al., 2014). Recently, the pentameric structure of the transmembrane domain within a lipid bilayer reconstituting the ERGIC membrane has been reported by NMR in the absence of the cytoplasmic part (Mandala et al., 2020). The C-terminal cytoplasmic part of protein E is important for interactions with different partners such as its PBM sequence to interact with PDZ-containing proteins.

**Figure 1.**
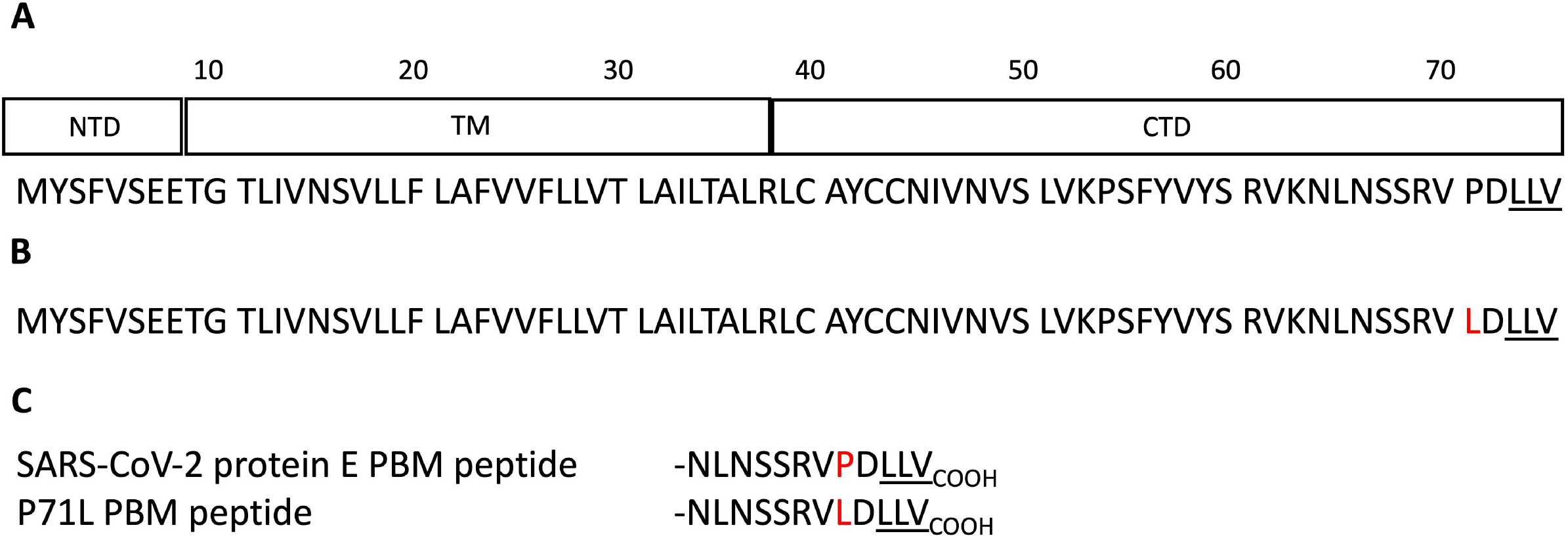
Schematic domain representation and sequences of the protein E constructs. (A) Domain delimitations and sequence of the full-length protein E of SARS-CoV-2. NTD, TM and CTD correspond to the N-terminal domain, the transmembrane domain and the C-terminal domain respectively. (B) Sequence of the full-length protein E of SARS-CoV-2 beta variant (B.1.351). (C) Sequences of the SARS-CoV-2 WT and the beta variant E C-terminal encompassing the PBM. The PBMs are underlined. The mutation is highlighted in red.

Previously, we used a high-throughput quantitative approach using a library covering all the human PDZ domains (Vincentelli et al., 2015) to establish the list of PDZ-containing proteins potentially targeted by SARS-CoV-2 E protein through its PBM. Among the ten PDZ-containing proteins identified, four are involved in cellular junction and polarity: ZO-1 (also called TJP1), LNX2, PARD3, MLLT4 (also called Afadin) (Caillet-Saguy et al., 2021)(Table 1).

**Table 1.**
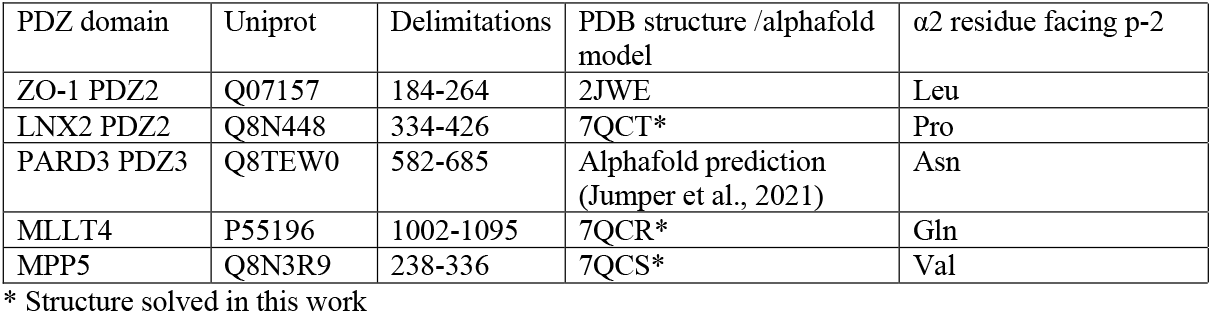
Selected human PDZ domains targeted by the SARS-CoV-2 E protein PBM. PDZ name, Uniprot code, delimitations of the constructs used in this work (residue numbers correspond to the numbering in the full-length human protein), PDB of the structure model used and the residue of PDZ helix α2 facing the PBM residue at position -2.

Indeed, ZO-1 is one of the essential proteins that connect transmembrane tight-junction proteins to the actin cytoskeleton. It consists of 3 PDZ domains, the second being the one that has a high affinity for the PBM motif of the E protein of SARS-CoV-2. *Robinot et al*. reported a disruption of the epithelial barrier integrity and an alteration of the ZO-1 distribution at tight junctions (TJs) during infection with SARS-CoV-2 (Robinot et al., 2021). LNX2 acts as a molecular scaffold for Numb family proteins, essential players in the regulation of cell adhesion and polarity (Wang et al., 2009). PARD3 is an essential protein in asymmetric cell division and in polarized growth. It plays a central role in the establishment of TJs (Chen et al., 2017). MLLT4 is an adapter protein linking nectin to the actin cytoskeleton, essential for the formation of adherent junctions and the regulation of cell adhesion (Ikeda et al., 1999). In addition, E protein of SARS-CoV was previously reported to interact with the PDZ-containing protein MPP5 (also called PALS1) through PBM-PDZ interaction altering TJ formation and the mammalian epithelium structure (Teoh et al., 2010). A recent study reported that the E protein PBM recognizes a pocket formed by residues from the PDZ and SH3 domains of MPP5 (Chai et al., 2021). As a key component of the Crumbs complex that controls the apical-basal polarity and TJ formation (Roh et al., 2002), MPP5 may contribute to the lung epithelium breakdown observed in patients infected by SARS-CoV. The relevance of these five PBM/PDZ interactions is not yet fully understood. Understanding the pathogenicity of this virus is currently a global health issue and targeting these virus-host interactions could reduce damage to the respiratory tract barrier and moderate the virus spread.

In this study, we showed that these five PDZ domains interact *in vitro* with the full-length E protein in a PBM-dependent manner and colocalize with the full-length E protein *in cellulo*, sequestrating the PDZ domains to the Golgi compartment. We further solved the three crystal structures of human PDZ/SARS-CoV-2 E PBM complexes for LNX2, MLLT4 and MPP5, highlighting their specific binding modes (Table 1). A mutation in E protein localized near the PBM (P71L) were reported in the variant of concern (VOC) beta (B.1.351)(Fig. 1B). We showed here different affinities between the PDZ domains and the SARS-CoV-2 WT and the beta variant E C-terminal encompassing the PBM (Fig. 1C). Thus, the acquired mutations might have important consequences on host machinery targeted during the infection in addition to a potential effect both on the structure and the ion-channel activity of the E protein.

## 2 Results

### 2.1 PDZ domains of the proteins involved in cell junctions and cell polarity bind to the full-length E protein in a PBM-dependent manner

We selected five PDZ-containing proteins involved in cellular junction and polarity from our previous high-throughput study on the specificity profile of the C-terminal SARS-CoV-2 E PBM sequence (Fig. 1C) against our library of all human PDZ domains (PDZome) using the automated holdup assay (Vincentelli et al., 2015; Duhoo et al., 2019; Caillet-Saguy et al., 2021). These are: ZO-1, LNX2, PARD3, MLLT4, and MPP5 previously reported to interact with SARS-CoV-2 E PBM through PBM-PDZ interactions (Table 1).

To investigate whether the five PDZ domains can interact with the full-length E protein and in a PBM-dependent manner, glutathione S-transferase (GST) pull-down assays were performed with the purified GST-tagged PDZ domains of ZO-1, LNX2, PARD3, MLLT4, and MPP5 and the lysates of HEK293 cells overexpressing constructs of GFP alone, GFP-tagged wild-type full-length E protein (GFP-E-WT), GFP-tagged full-length E protein with the PBM mutated with glycines (GFP-E-GGGG) and GFP-tagged cytoplasmic tail of protein E (last 12 residues) (GFP-E last 12 aa) designated 1, 2, 3 and 4 in figure 2 (Fig. 2A). The GFP-tagged constructs expression was assessed by examining the fluorescence emitted by the GFP by microscopy and by Western-Blot on the cell lysates using anti-GFP antibody (Fig. 2B). GST alone was used as a negative control. We confirmed using Ponceau S staining that equal amount of GST-tagged protein constructs was bound to the GST resin (Fig. 2C). GST alone and its fusion with PDZ ZO1, PDZ LNX2, PDZ MLLT4, PDZ MPP5 and PDZ PARD3, are used as baits and were immobilized on glutathione beads and tested for their ability to pull down GFP alone, GFP-E-WT, GFP-E-GGGG and GFP-E last 12 aa by Western-blot using the anti-GFP antibody. After washing, identical amounts of beads were analysed for the presence of GFP-tagged proteins.

**Figure 2.**
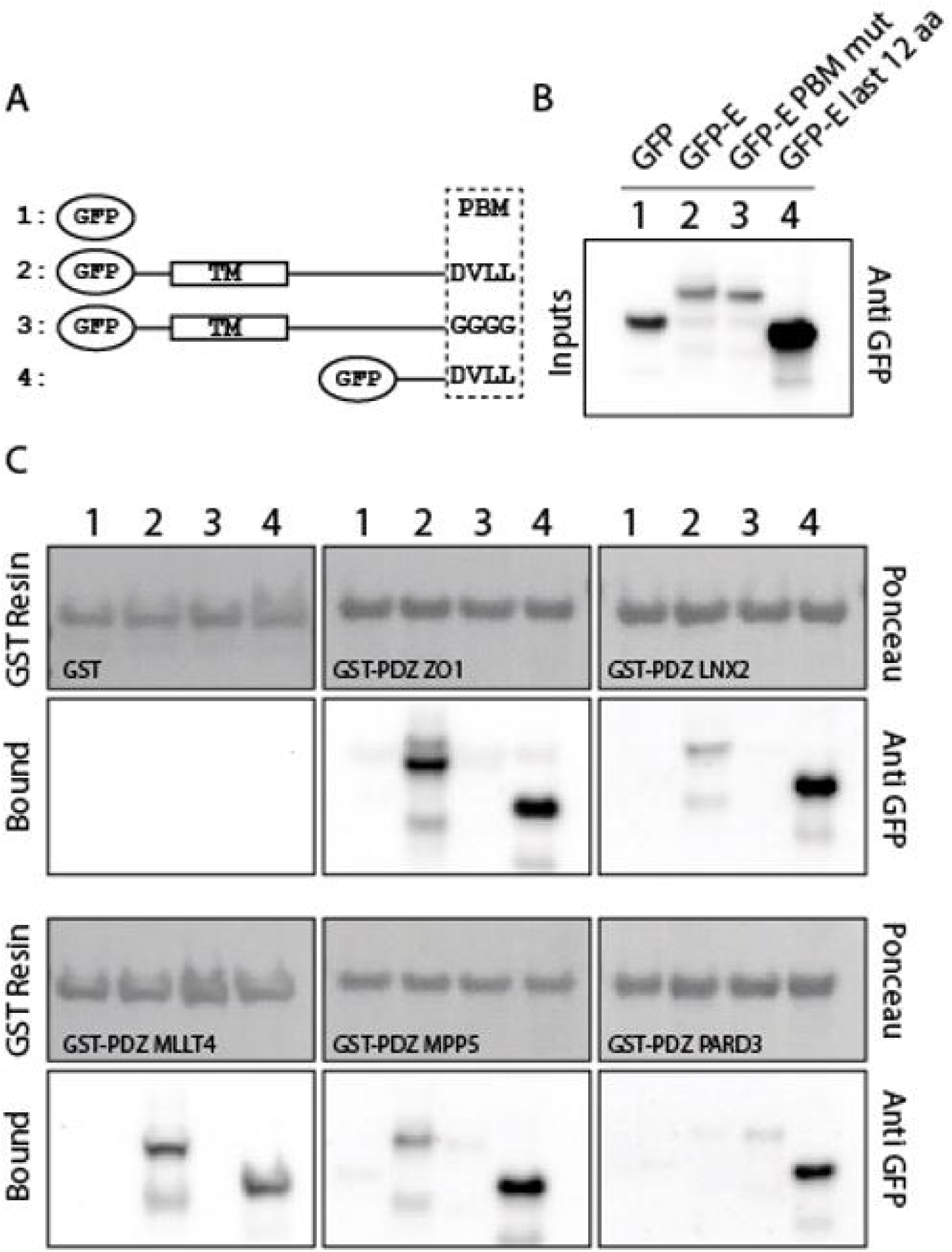
PDZ domains interact with the SARS-CoV-2 full-length E protein in a PBM-dependent manner. (A) Schematic representation of the GFP-E constructs used in this study. TM and PBM correspond to the transmembrane domain and to the PDZ-Binding Motif respectively. (B) Input fraction of GFP tagged viral E gene construct expressed in HEK293 cells and used for pull-down. Samples were analysed by immunoblotting using anti-GFP antibody. (C) GST pull-down binding results with immobilised GST or GST-PDZ domains (GST-PDZ ZO1, GST-PDZ LNX2, GST-PDZ MLLT4, GST-PDZ MPP5 and GST-PDZ PARD3) used as affinity resin and incubated with the HEK293 cell lysates. GST-tagged proteins were stained by Ponceau S (top panels). The bound fraction was analysed by immunoblotting using an anti-GFP antibody (bottom panels). Note that blots represent discontinuous panels from the gels when black line delimitation is present. The images have been cropped to frame the relevant region.

The GFP-E last 12 aa was detected in all interactions with GST-PDZ domains but not detected with GST alone confirming the interactions identified in our high-throughput holdup assay (Caillet-Saguy et al., 2021) such validating the interactions within the context of lysates of HEK293 cells overexpressing SARS-CoV-2 protein E constructs (Fig. 2C).

GFP-E-WT was also detected in all interactions with GST-PDZ domains but with GST-PDZ PARD3 that showed no clear band (Fig. 2C). In all cases, the bands have a weaker intensity than GFP-E last 12 aa in agreement with a significant lower expression in cells, as shown in the inputs for GFP-E-WT compared to GFP-E last 12 aa (Fig. 2B). We failed to obtain a clear detection of the band for GST-PDZ PARD3 (Fig 2C). Conversely, GFP-E-GGGG was not detected in all interactions with GST-PDZ domains except with GST-PDZ PARD3 that showed a weak band (Fig. 2C).

Altogether, these results indicate a specific interaction between the SARS-CoV-2 WT protein E and the PDZ domains of ZO-1, LNX2, MLLT4 and MPP5 with variations in binding intensities except for PARD3. These interactions are PBM-dependent in the context of the full-length E protein.

### 2.2 Structures of complexes between PDZ domains of MLLT4, MPP5 and LNX2 and the C-terminal of protein E

We deciphered the molecular basis of recognition of the C-terminal sequence of the protein E of SARS-CoV-2 encompassing the PBM by the PDZ domains of the proteins engaged in cellular junction and polarity. To this aim, we solved the crystal structures of the complex formed by the C-terminal peptide of protein E encompassing the PBM (Fig 1C) and the PDZ domains of MLLT4, MPP5 and LNX2 (Table 1) by molecular replacement (Fig. 3; Table 2). We have described the set of intermolecular bonds to gain structural insights into the binding mode of the C-terminal sequence of the protein E of SARS-CoV-2 with PDZ domains.

**Figure 3.**
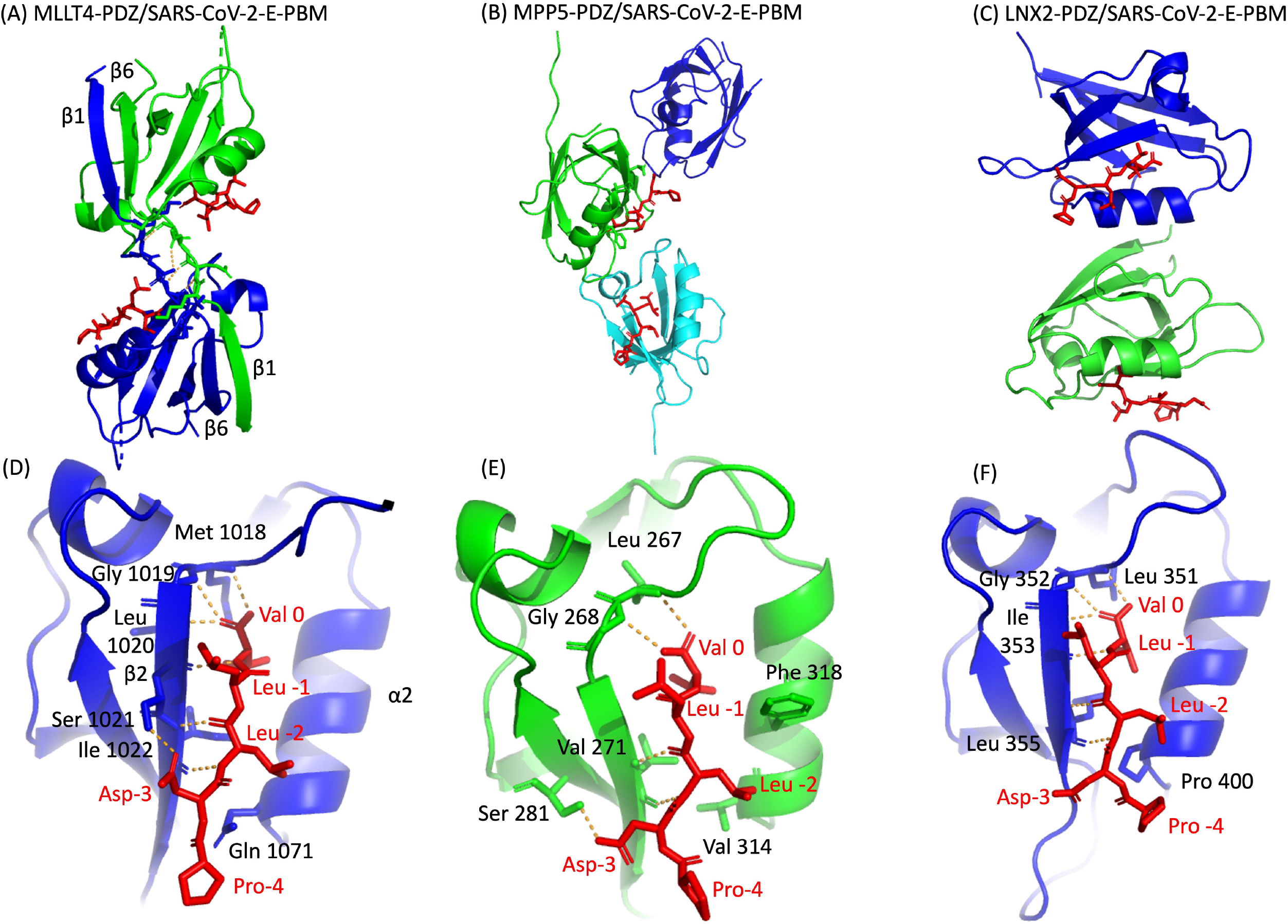
X-ray structures of PDZ domains of MLLT4, MPP5 and LNX2 bound to the SARS-CoV-2 protein E PBM. (A)(B)(C) The asymmetric unit of the PDZ domains of MLLT4, MPP5 and LNX2 respectively, bound to the SARS-CoV-2 protein E PBM shown as red sticks. (A) Selected interchain polar contacts related to the swapped dimer between the fragments Lys 1014-Gly 1017 of each chain are shown in orange and the associated residues are shown as sticks. (D)(E)(F) Detailed views of the PDZ domains bound to SARS-CoV-2 protein E PBM. Important residues are labeled and shown as sticks. Intermolecular H-bonds and polar contacts are reported as orange dashed lines.

**Table 2.**
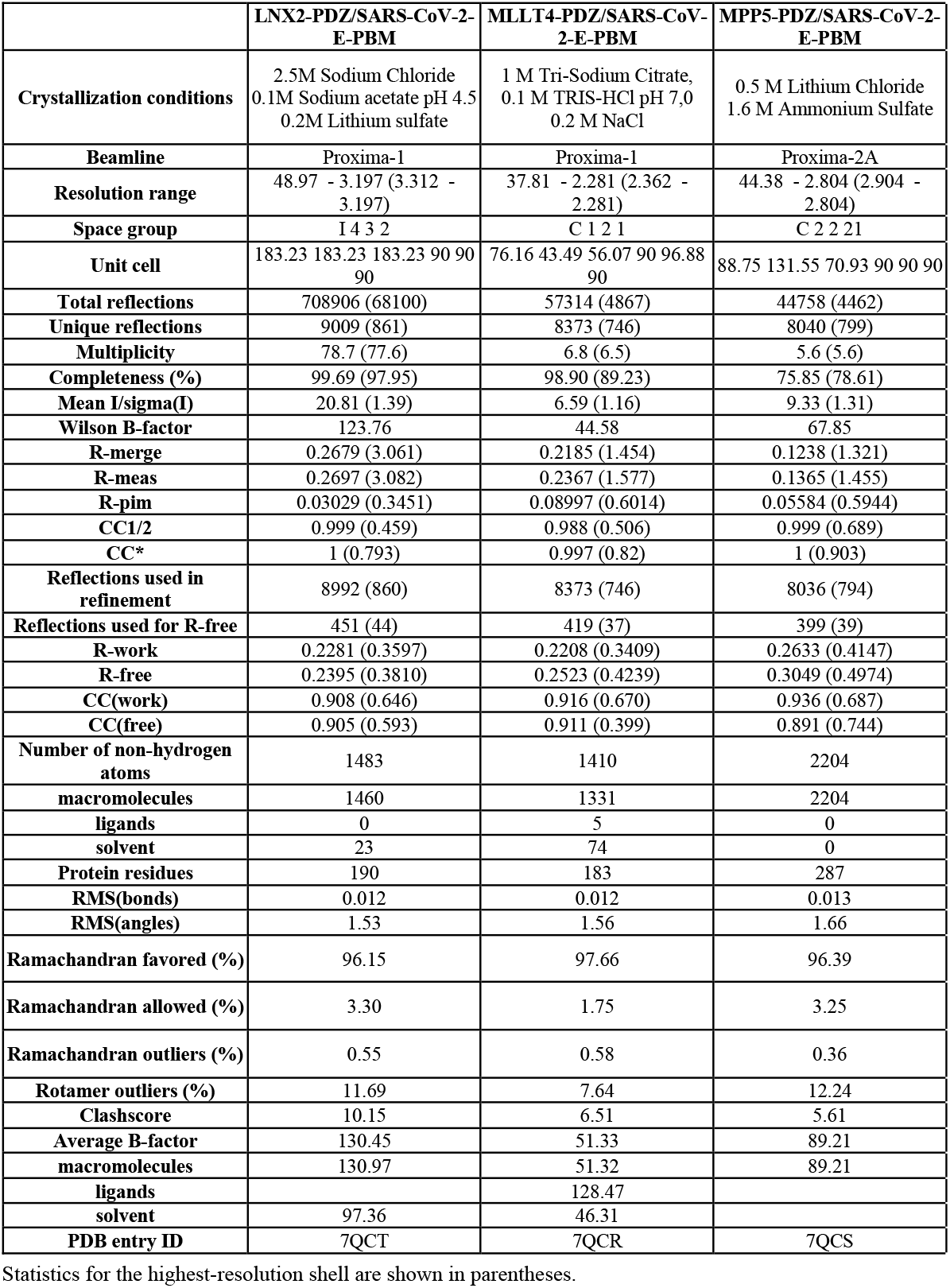
Data collection and refinement statistics.

|  | **LNX2-PDZ/SARS-CoV-2-E-PBM** | **MLLT4-PDZ/SARS-CoV-2-E-PBM** | **MPP5-PDZ/SARS-CoV-2-E-PBM** |
| --- | --- | --- | --- |
| **Crystallization conditions** | 2.5M Sodium Chloride<br>0.1M Sodium acetate pH 4.5<br>0.2M Lithium sulfate | 1 M Tri-Sodium Citrate,<br>0.1 M TRIS-HCl pH 7.0<br>0.2 M NaCl | 0.5 M Lithium Chloride<br>1.6 M Ammonium Sulfate |
| **Beamline** | Proxima-1 | Proxima-1 | Proxima-2A |
| **Resolution range** | 48.97 - 3.197 (3.312 - 3.197) | 37.81 - 2.281 (2.362 - 2.281) | 44.38 - 2.804 (2.904 - 2.804) |
| **Space group** | I 4 3 2 | C 1 2 1 | C 2 2 21 |
| **Unit cell** | 183.23 183.23 183.23 90 90 90 | 76.16 43.49 56.07 90 96.88 90 | 88.75 131.55 70.93 90 90 90 |
| **Total reflections** | 708906 (68100) | 57314 (4867) | 44758 (4462) |
| **Unique reflections** | 9009 (861) | 8373 (746) | 8040 (799) |
| **Multiplicity** | 78.7 (77.6) | 6.8 (6.5) | 5.6 (5.6) |
| **Completeness (%)** | 99.69 (97.95) | 98.90 (89.23) | 75.85 (78.61) |
| **Mean I/sigma(I)** | 20.81 (1.39) | 6.59 (1.16) | 9.33 (1.31) |
| **Wilson B-factor** | 123.76 | 44.58 | 67.85 |
| **R-merge** | 0.2679 (3.061) | 0.2185 (1.454) | 0.1238 (1.321) |
| **R-meas** | 0.2697 (3.082) | 0.2367 (1.577) | 0.1365 (1.455) |
| **R-pim** | 0.03029 (0.3451) | 0.08997 (0.6014) | 0.05584 (0.5944) |
| **CC1/2** | 0.999 (0.459) | 0.988 (0.506) | 0.999 (0.689) |
| **CC\*** | 1 (0.793) | 0.997 (0.82) | 1 (0.903) |
| **Reflections used in refinement** | 8992 (860) | 8373 (746) | 8036 (794) |
| **Reflections used for R-free** | 451 (44) | 419 (37) | 399 (39) |
| **R-work** | 0.2281 (0.3597) | 0.2208 (0.3409) | 0.2633 (0.4147) |
| **R-free** | 0.2395 (0.3810) | 0.2523 (0.4239) | 0.3049 (0.4974) |
| **CC(work)** | 0.908 (0.646) | 0.916 (0.670) | 0.936 (0.687) |
| **CC(free)** | 0.905 (0.593) | 0.911 (0.399) | 0.891 (0.744) |
| **Number of non-hydrogen atoms** | 1483 | 1410 | 2204 |
| **macromolecules** | 1460 | 1331 | 2204 |
| **ligands** | 0 | 5 | 0 |
| **solvent** | 23 | 74 | 0 |
| **Protein residues** | 190 | 183 | 287 |
| **RMS(bonds)** | 0.012 | 0.012 | 0.013 |
| **RMS(angles)** | 1.53 | 1.56 | 1.66 |
| **Ramachandran favored (%)** | 96.15 | 97.66 | 96.39 |
| **Ramachandran allowed (%)** | 3.30 | 1.75 | 3.25 |
| **Ramachandran outliers (%)** | 0.55 | 0.58 | 0.36 |
| **Rotamer outliers (%)** | 11.69 | 7.64 | 12.24 |
| **Clashscore** | 10.15 | 6.51 | 5.61 |
| **Average B-factor** | 130.45 | 51.33 | 89.21 |
| **macromolecules** | 130.97 | 51.32 | 89.21 |
| **ligands** |  | 128.47 |  |
| **solvent** | 97.36 | 46.31 |  |
| **PDB entry ID** | 7QCT | 7QCR | 7QCS |

716 Statistics for the highest-resolution shell are shown in parentheses.

#### 2.2.1 Structure of the PDZ domain of MLLT4 in complex with the PBM of protein E

The crystal structure of MLLT4-PDZ in complex with SARS-CoV-2-E-PBM peptide was solved at a resolution of 2.28 Å. The final refined model contains two PDZ domains per asymmetric unit that adopt a swapped dimer conformation with the PDZ folding comprising five β strands and two α helices. Two peptides are bound to each PDZ dimer (Fig. 3A). An electron density is observed for the last five and last three residues of the peptide containing the PBM indicating a well-defined conformation of these last C-terminal residues. The last three residues of the SARS-CoV-2-E-PBM peptide bind to the α2/β2-groove in each PDZ unit (Fig. 3D).

The swapped dimer comprises an intermolecular interaction between the two PDZ domains through the β1/β6 pair (Fig. 3A), and several pairs of inter-domain H-bonds between the fragments Lys 1014-Gly 1017 of each chain (Fig. 3A).

The E PBM binds to the PDZ in a conventional manner as an antiparallel extension of the β2 strand by inserting into a binding groove formed by the β2 strand, the α2 helix and the “GLGF” motif. The last four residues of the SARS-CoV-2-E-PBM peptide were in contact with the PDZ domain, whereas only the last three residues were in contact and the upstream residues were distant from the PDZ domain surface in the previously reported complex between MLLT4-PDZ with the nectin-3 or the Bcr PBM peptides (Chen et al., 2007; Fujiwara et al., 2015).

The C-terminal carboxylate of Val at position 0 of the PBM forms three hydrogen bonds with the amide protons of Met 1018, Gly 1019 and Leu 1020 of the “GLGF” loop of MLLT4-PDZ (Fig. 3D). A hydrogen bond is also formed between the proton amide of this Val 0 and the carbonyl of Leu 1020 of the β2 strand. In addition, the carbonyl and the amide group of Leu at position -2 interacts through a hydrogen bond with the proton amide and the carbonyl group of Ile 1022, respectively. The Ile -2 side chain establish hydrophobic contacts with the side chains of Gln 1071 and Ile 1022. A hydrogen bond is also formed between the carboxyl at the end of the side chain of Asp at position - 3 of the PBM and the hydroxyl of Ser 1021 of the β2 strand (Fig. 3D). Thus, the formed β sheet involved the last three residues of the SARS-CoV-2 E protein.

#### 2.2.2 Structure of the PDZ domain of MPP5 in complex with the PBM of protein E

The crystal structure of the MPP5-PDZ in complex with SARS-CoV-2-E-PBM was solved at a resolution of 2.80 Å. There are three MPP5-PDZ per asymmetric unit (Fig. 3B). Each domain adopts a compact globular PDZ fold, and an electron density was observed for the last five residues of the peptide in two MPP5-PDZ present in the asymmetric unit.

As for MLLT4, the E PBM conventionally binds to the MPP5-PDZ. The C-terminal carboxylate of the Val at position 0 of the PBM forms two hydrogen bonds: one with the amide proton of Leu 267 and one with Gly 268 of the “GLGF” loop of MPP5-PDZ (Fig. 3E). In addition, the amide and carbonyl groups of Leu at position -2 form hydrogen bonds with the carbonyl and amide proton of valine 271, respectively. Interestingly, as for MLLT4, a hydrogen bond is also formed between the carboxyl at the end of the side chain of Asp at position - 3 of the PBM and the hydroxyl group of Ser 281 of the β3 strand (Fig. 3E). The key hydrogen bonds of Val 0 and Leu -2 are similar with the structure recently reported (Javorsky et al., 2021). However, the side chains of Asp at position -3 and Arg 272 form an ionic bond within this structure. The alternative hydrogen bond with Ser 281 identified in our structure was previously reported in the complex formed by MPP5-PDZ and the PBM peptide of Crumbs (Javorsky et al., 2021). Furthermore, the side chain of Phe 318 in α2 helix, which prevents access to the binding groove in the unbound form (Ivanova et al., 2015), is located out of the binding groove in our structure allowing SARS-CoV-2-E-PBM peptide binding (Fig. 3E) as previously stated for MPP5-PDZ in complex with SARS-CoV-1 E and SARS-CoV-2 E PBMs, as well as for Crumbs peptides (Javorsky et al., 2021).

#### 2.2.3 Structure of the PDZ domain of LNX2 in complex with the PBM of protein E

The crystal structure of LNX2-PDZ2 in complex with SARS-CoV-2-E-PBM was solved at a resolution of 3.3 Å. Each of the two LNX2-PDZ present in the asymmetric unit are bound to the peptide (Fig. 3C). The PDZ fold agrees with the unbound form of LNX2-PDZ (PDB 5e1y) with an overall RMSD of the backbone atoms of the two PDZ of 0.44 Å. Electron density was observed for the last five residues of the two peptides (Fig. 3F). In both cases, the last three residues of the SARS-CoV-2-E-PBM peptide bind to the canonical α2 helix /β2 strand groove in each PDZ unit (Fig. 3C).

As for MLLT4 and MPP5, the PBM of protein E conventionally binds to the PDZ of LNX2. The C-terminal carboxylate of Val 0 of the PBM forms three hydrogen bonds: two with the amide protons of Leu 351 and Gly 352 of the “GLGF” loop of LNX2-PDZ2 and one with the Ile 353 amide proton (Fig. 3F). The proton amide of Val 0 forms also a H-bond with the carbonyl of Ile 353. In addition, the amide and carbonyl groups of Leu -2 form hydrogen bonds with the carbonyl and amide proton of Leu 355 respectively (Fig. 3F). In one of the two complexes, an ionic bond is formed between the carboxylate of Val at position - 3 of the PBM and the amine group of Arg 357 at the end of the β2 strand.

In summary, we have shown that the SARS-CoV-2-E PBM peptide interacts with the PDZ domains of MLLT4, MPP5 and LNX2 with similar binding modes at position 0 and -2. The SARS-CoV-2-E PBM is a class II PBM with a hydrophobic residue at position -2 that should contact a hydrophobic residue or the aliphatic part of a Lysine at the N-terminal of α2 helix of the PDZ domain (Songyang et al., 1997; Harris and Lim, 2001). Indeed, the N-terminal of α2 helix is occupied by a valine (Val 314), a proline (Pro 400), and a glutamine (Gln 1071) in MPP5-PDZ, LNX2-PDZ and MLLT4-PDZ, respectively. A glutamine at this position is different from a canonical class II PDZ but still classified as a class II PDZ (Fig. 3; Table 1). The side chain of aspartic acid at position -3 is involved in H-bond with MLLT4-PDZ and MPP5-PDZ whereas the proline at position -4 does not interact with none of the PDZ domains. Thus, we established the structural basis of SARS-CoV-2-E PBM binding to the PDZ domains of MLLT4, MPP5 and LNX2.

### 2.3 The viral E protein sequestrates PDZ domains to the Golgi compartment

Then we explore the PDZ-PBM interactions between the full-length E protein and the selected PDZ domains in cells.

SARS-CoV-1 or SARS-CoV-2 E proteins expressed from cDNA were reported to localize to the endoplasmic reticulum (ER), ERGIC, or Golgi complex depending on the nature and localization of the tag (N or C) was used in the study (Lisa A. Lopez, 2005; Cohen et al., 2011; Pearson et al., 2021). Here we used the recently developed ALFA tag, that is small (14 residues) and electroneutral (Götzke et al., 2019) to tag the SARS-CoV-2 E protein at the N-terminal. HeLa cells transiently transfected with SARS-CoV-2 ALFA-E encoding plasmids displayed strong Golgi expression of the viral protein as revealed by the co-staining with Golgi marker GM130 (Supplementary Figure S1A).

We then investigated the ability of the viral E protein to recruit GFP-tagged PDZ domains to the Golgi compartment. When transfected alone, GFP-tagged PDZ domains from ZO1, MLLT4, MPP5/PALS1, LNX2 and PARD3 do not accumulate in the Golgi apparatus (supplementary Figure S1B). Interestingly, all these GFP-tagged PDZ domains relocalize to the Golgi compartment when they are co-transfected with the ALFA-E construct as shown by the GM130 co-staining for ZO1 (Fig. 4A) and the other PDZ domains (Supplementary Figure S1C). ZO-1 PDZ2/E protein colocalization is strictly PBM-dependent as no colocalization is observed between ZO1-PDZ2 and the ALFA E PBM mutant for which the PBM sequence DVLL was substituted by four glycines (Fig. 4B). Likewise, this interaction was specific as SCRIB-PDZ1 domain, an unrelated PDZ domain that was not identified in our screen, showed no tropism for the Golgi compartment when it was cotransfected with the ALFA E construct (Fig. 4C). These results indicate that the viral E protein can bind to ZO1, MLLT4, MPP5, LNX2 and PARD3 PDZ domains *in cellulo* in a PBM specific manner and recruit them to the Golgi apparatus.

**Figure 4.**
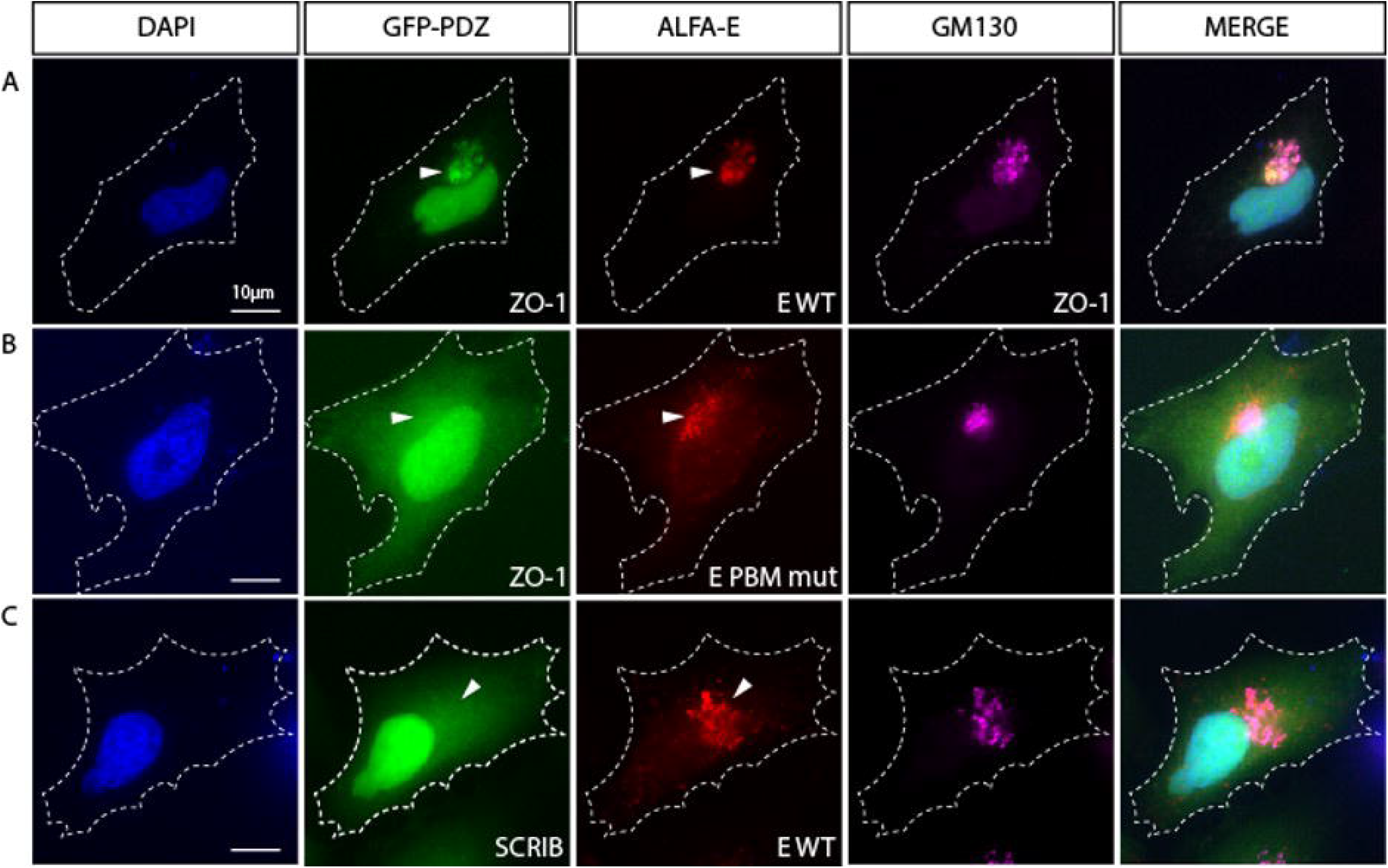
The viral E protein recruits the PDZ domain of ZO1 to the Golgi apparatus. (A) Hela cell co-transfected with encoding constructs GFP-PDZ2 of ZO1 and ALFA-E (red) displays Golgi recruitment of both proteins as indicated by GM130 staining (Magenta). (B) Co-transfection of GFP-ZO1 with ALFA-E PBM mutant (red) does not lead to the recruitment of GFP-ZO1 to the Golgi apparatus. (C) Hela cell co-transfected with GFP-PDZ1 of SCRIB and ALFA-E (red) does not show recruitment of the proteins to the Golgi apparatus (Magenta). White arrowheads indicate Golgi apparatus position. Nuclei are stained with DAPI. Bars correspond to 10µm.

### 2.4 The PDZ domains bind with different affinities the original C-terminal SARS-CoV-2-E PBM and the C-terminal SARS-CoV-2-E PBM P71L mutant of the VOC beta

Acquired mutations in SARS-CoV-2 protein E localized close to the PBM were reported and the most common non-synonymous mutations were S68F and P71L (Hassan et al., 2020). We focused on the P71L mutation found in the beta variant. The VOC beta was first identified in South Africa in September 2020 and has been reported in more than 130 countries. The mutations on the beta variant make it more transmissible (Tegally et al., 2021), with greater antibody resistance compared to earlier variants of SARS-CoV-2. The beta variant is estimated to be 50% more infectious than the original coronavirus strain (Tegally et al., 2021), and could be of a higher risk of hospitalization, admission to intensive care, and death (Funk et al., 2021).

The P71L mutation in the E protein of the beta variant is in the vicinity of the PBM at position -4. Upstream residues can also affect the PDZ binding affinity and specificity in addition to the C-terminal positions 0 and -2 of the PBM, which are important for canonical PDZ domain binding (Luck et al., 2012).

We examined the binding affinities of the C-terminal SARS-CoV-2-E PBM peptide (12-mers peptide NLNSSRVPDLLV_COOH_) from the original strain and the P71L PBM peptide (12-mers peptide sequence NLNSSRVLDLLV_COOH_) against the PDZ domains of ZO1, MLLT4, MPP5, LNX2 and PARD3 using microscale thermophoresis (MST)(Table 3; Supplementary Figure S2). The averaged affinity constants (Kds) of the two PBM peptides for the different PDZ partners are summarized in Table 3. The best affinities for the original SARS-CoV-2 peptide E are obtained with the PDZ2 of ZO-1 and the PDZ of MPP5 with Kd values of 15 μM and 30 μM, respectively. The affinities of SARS-CoV-2 peptide E for PARD3-PDZ3 and MLLT4-PDZ are lower with Kd values of 341 µM and 569 µM, respectively. The beta variant P71L mutation does not significantly impact the affinity for the PDZ of MPP5 but considerably reduces the one for the ZO-1-PDZ2 with a Kd of 133 µM, and the PDZ domains of MLLT4 and PARD3 with affinities not detectable in the tested concentration range. Thus, either the P71L mutant does not interact with the PDZs of MLLT4 and PARD3, or the affinities are outside the tested concentration range (Kd> 1 mM).

**Table 3.**
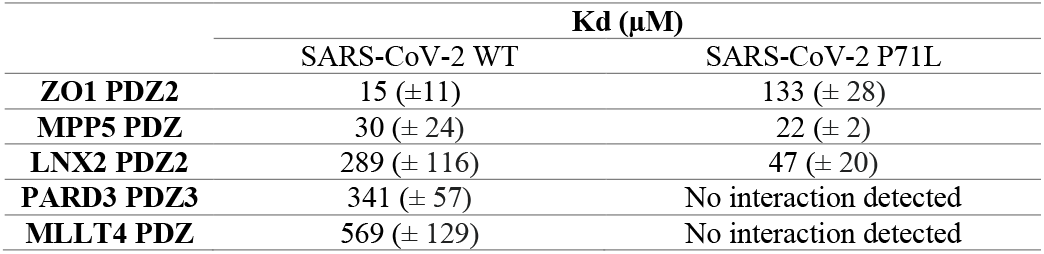
Kd values between the E protein PBM of SARS-CoV-2 WT and of the P71L mutant for the selected PDZ domains. The data are representative of two independent experiments and error bars correspond to the standard deviation.

Conversely, the affinity of LNX2-PDZ2 is improved from 289 µM to 47 µM with the mutant P71L. Thus, this mutation leads to a significant increase in the affinity for LNX2-PDZ2 with a Kd almost 6 times lower compared to the one of SARS-CoV-2 WT.

To resume, significant interactions are detected between the original C-terminal SARS-CoV-2-E PBM peptide and all the PDZ domains we tested by MST, in agreement with our previous results using the high-throughput screening Holdup assay (Caillet-Saguy et al., 2021). Interestingly, we observed different binding affinities for four over five PDZ domains with the P71L mutant. This strongly suggests a differential specificity profile against the PDZome between the original C-terminal SARS-CoV-2-E PBM and the C-terminal SARS-CoV-2-E PBM P71L mutant.

## 3 Discussion

Deletion of protein E strongly reduces the replication of SARS-CoV-1 in cells and suppresses virus induced mortality in mice (DeDiego et al., 2007). The protein E of SARS-CoV-1 contains a C-terminal type II PBM involved in viral pathogenesis (Jimenez-Guardeño et al., 2014) that binds to PDZ domains of cellular proteins. Interestingly, mice infected with a E PBM mutant virus showed 100% survival rate, mimicking the phenotype seen with the deletion of the full length E protein (Jimenez-Guardeño et al., 2014). The protein E from SARS-CoV-2 presents also a PBM like that of SARS-CoV-1 and we recently identified the PDZ proteins targeted by the PBM of SARS-CoV-1, SARS-CoV-2 and MERS by screening *in vitro* the full human PDZ library (Caillet-Saguy et al., 2021). From this previous study and others (Teoh et al., 2010; Toto et al., 2020), we selected five human PDZ-containing proteins showing significant interactions with the SARS-CoV-2 protein E PBM and involved in cellular junctions and cell polarity: ZO-1/TJP1, PARD3, MLLT4, LNX2 and MPP5/Pals1. Indeed, targeting cellular junctions and polarity machineries is a shared strategy by viruses to facilitate the infectious cycle improving either viral entry, replication, dissemination, or egress. Notably, ZO-1 and Pals1 are targeted by other viruses such as influenza and SARS-CoV, to disrupt and open TJs to efficiently exit the airway epithelia to spread and disseminate (Torres-Flores and Arias, 2015). SARS-CoV-2 was shown to transiently impair bronchial epithelium altering the distribution of ZO-1 (Robinot et al., 2021) and Pals1 was shown to translocate from TJ to ERGIC when is targeted by protein E through its PBM resulting in the dissociation of TJ in epithelia of various organs (Chai et al., 2021). In this study, we showed that the five PDZ domains TJP1, PARD3, MLLT4, LNX2 and MPP5/PALS1 interact in a PBM-dependent manner *in vitro* and these five PDZ domains colocalize with the full-length E protein *in cellulo* sequestrating the PDZ domains into the Golgi compartment. This is consistent with the exposition of the C-terminus of the protein E to the cytoplasmic side as previously reported (Duart et al., 2020), allowing interactions with viral and host proteins.

Interestingly, our pull-down experiments showed that the binding partners identified in the holdup screening can interact not only with the last 12 amino acids, as previously shown in the holdup screening, but also with the full-length E protein. We noted a weaker detection of the interaction with the full-length WT protein E compared to the C-terminal peptide because the protein is less expressed, formed oligomers and the C-terminal peptide is likely less accessible in the context of the entire protein inserted in the membrane (Park et al., 2021). Moreover, the overexpressed protein E induced a significant cellular mortality. It was previously proposed that the cation channel formed by the overexpressed protein E could disrupt the host cell membrane (Cao et al., 2021). Less cells are recovered after transfection with WT protein E than with other constructs. To address this issue, we transfected more cells. Interestingly, no cellular toxicity was observed with GFP-E-GGGG construct. This is in agreement with a previous work on SARS-CoV-1 that showed that the mutant E-GGGG is no longer inducing lethality in mice (Jimenez-Guardeño et al., 2014), illustrating the impact of the E PBM on the homeostasis of infected cells.

Altogether, our results indicate significant interactions between the SARS-CoV-2 WT protein E and the PDZ domains of ZO-1, LNX2, MLLT4 and MPP5. These interactions are dependent on the presence of the PBM in the full-length protein E since the mutation of the PBM led to a loss of interaction. PARD3 binds the SARS-CoV-2 E PBM *in vitro* and colocalizes with the full-length protein E and not with the mutated PBM E protein that also strongly suggest a specific interaction. We solved the X-ray structures of the complexes between the SARS-CoV-2 E C-terminal encompassing the PBM, and the human LNX2, MLLT4 and MPP5/Pals1 PDZs highlighting the binding modes for three of the potential cellular targets of protein E. We found a swapped dimer in the structure of MLLT4-PDZ with the β1 and β6 strands swapped (PDB 7QCR). Previous NMR structures of MLLT4-PDZ complexed or not with other compounds (peptides or small molecules) only reported monomers (PDB 1XZ9, 2EXG, 1T2M, 2AIN). One X-ray structure reported a dimer due to a fusion of a PBM at the C-terminus of MLLT4-PDZ that created a new dimer interface forming an antiparallel β sheet between β2 strands (PDB 3AXA). To investigate the presence of a dimer in solution, we performed Analytical Ultracentrifugation (AUC) experiments on MLLT4-PDZ (Table 4; Supplementary Figure S3). The MLLT4-PDZ is found mainly monomeric in solution with a sedimentation coefficient of 1.4S, suggesting that the swapped dimer is probably an artefact of crystallization. Interestingly, MLLT4-PDZ is able to recognize type II PBM but also type I PBM due to an unexpected glutamine in the α2 helix as previously reported (Zhou et al., 2005).

**Table 4.** Hydrodynamic parameters of ZO-1 PDZ2 and MLLT4 PDZ derived from the analysis of analytical ultracentrifugation.

|  | ZO-1 1 mg/ml |  |  | MLLT4-PBM 1mg/ml |  |  |
| --- | --- | --- | --- | --- | --- | --- |
| Sedimentation coefficient (S) | 2.1 | 2.8 | 3.7 | 0.5 | 1.4 | 2.3 |
| Frictional ratio $f/f_0$ (average) | 1.4 | | | 1.2 | | |
| Molecular weight estimated by AUC (kDa) | 26 | 40 | 60 | 2.8 | 12 | 26 |
| Peak proportion % | 49 | 49 | 2 | 25 | 65 | 10 |

Remarkably, ZO-1-PDZ2 forms a very stable swapped dimer with an extended antiparallel inter-domain β sheet. The β1 and β2 strands of one domain is swapped with those from the second domain allowing the formation of the binding groove (Fanning et al., 2007; Chen et al., 2008). We also performed AUC experiments on ZO-1-PDZ2 to verify its oligomeric state in solution (Table 4; Supplementary Figure S3). We found that ZO-1-PDZ2 adopts two oligomeric states in solution: one dimeric form and one higher oligomeric state possibly compatible with a tetramer in solution that might be consistent with a swapped dimer in solution. Conclusively, the X-ray structures have provided the structural basis for SARS-CoV-2-E PBM binding to the PDZ domains of MLLT4, MPP5 and LNX2 and these results offer a mechanistic beginning for SARS-CoV-2 perturbation of PDZ-containing proteins.

Finally, we showed different affinities for the cellular PDZ domains of ZO-1, LNX2, PARD3, MLLT4 and MPP5, with the original SARS-CoV-2 E C-terminal peptide (-NLNSSRVPDLLV_COOH_) and the beta variant E C-terminal peptide (-NLNSSRVLDLLV_COOH_) containing the P71L mutation close to the PBM (Fig. 1B). This position –4 is not well defined within an electronic density in the crystal structures and when a density is observed, it is the last upstream residue modeled for the C-terminal of the protein E. In the three complexes we studied, the residue in position -4 is not involved in the interaction network with the PDZ domains. Surprisingly, we determined that P71L mutation markedly altered affinities with host PDZ domains. While it noticeably reduces the affinities for the PDZ domains of ZO-1, MLLT4 and PARD3, it enhances its affinity for LNX2 compared to SARS-CoV-2 WT. Influences of PBM upstream residues on the affinity for PDZ domains has already been reported previously (Lee and Zheng, 2010; Terrien et al., 2012). These findings show that the acquired P71L mutation might have important effects on the human PDZome targeted by the variant during the infection.

Less prevalent than the alpha and delta variants, the beta variant accounted for approximately 10% of virus samples in France in June 2021 (https://www.santepubliquefrance.fr/). Mutations in the beta variant make it more transmissible, with greater antibody resistance, higher risk of hospitalization, ICU admission, and death compared to earlier SARS-CoV-2 E variants (Veneti et al., 2021; Wang et al., 2021). In addition to the P71L mutation, the beta variant has other mutations, including three on its spike protein, which may help the virus to escape antibodies and to bind more tightly to human cells (Han et al., 2021). The importance of the P71L mutation in the greater pathogenicity of the beta variant remains to be demonstrated. Nevertheless, because viroporins like E protein are linked to inflammasome activation during viral infection (Schoeman and Fielding, 2019), mutations that can alter the specificity of E for its protein partners could potentially impact the viral E pro-inflammatory behavior and thus be associated with a change in viral pathogenicity.

## 4 Material and methods

### 4.1 Constructs cloning

DNA sequences encoding the PDZ domains of ZO-1-2 (PDZ2; residues 184-264), LNX2-2 (PDZ2; residues 334-426), PARDD3-3 (PDZ3; residues 582-685), MLLT4 (residues 961-1056) and MPP5 (residues 238-336) come from the PDZome library with PDZ domains cloned into a Gateway system (Vincentelli et al., 2015; Duhoo et al., 2019). The PDZ domains were cloned into pETG-41A plasmid vectors as an N-terminal fusion to a histidine tag and a maltose-binding protein (His-MBP-tag). Optimization of expression and purification conditions led us to test other tags for most constructs using the Gateway vector recombination system. Following the manufacturer protocol of the Gateway™ LR Clonase™ II Enzyme mix (Invitrogen), PDZ domains of ZO-1, LNX2 and PARD3 were subcloned into pDEST™17 vector and MPP5-PDZ into pDEST™15 vector, allowing the production of recombinant proteins as a fusion to an N-terminal histidine and GST tag, respectively. A TEV cleavage site was introduced between the N-terminal tags and the PDZ sequences.

The pCMV ALFA E vector was constructed as follows: DNA sequence encoding the ALFA tag (MSRLEEELRRRLTE) followed by a linker (GGGGS) fused to the sequence corresponding to the alpha-variant of SARS-CoV2 E cDNA (GenBank: BCM16077.1) synthetized by Eurofins. The ALFA-linker-E sequence was subsequently cloned into a pCMV backbone vector using ClaI and XhoI. The E PBM C-terminal mutation (DVLL>GGGG) was performed using the Q5 site-directed mutagenesis kit (NEB). The sequences corresponding to the PDZ domains were cloned into the pCMV GFP vector using EcoRI and XhoI sites.

### 4.2 Protein expression and purification

The vectors were used to transform *E. coli* BL21 Star (DE3) (Invitrogen) strain. Bacteria were grown in LB medium supplemented with ampicillin (100 mg / L) at 37°C. Protein expression was induced at OD600nm 0.8-1 with 0.2 mM IPTG at 18°C overnight. Bacteria were then harvested and PDZ domains of MLLT4, LNX2, PARD3 and MPP5 were resuspended in lysis buffer (Tris 50 mM (pH 7.5), NaCl 250 mM, β-mercaptoethanol 2 mM, protease inhibitors, one tablet per 50 mL of buffer (EDTA-free, Roche Diagnostics) and benzonase (E1014-25KU >250 Units/liter of culture)). Then, the cells were broken under pressure by CELLD (1.3 kBar at 4°C). The debris were pelleted by centrifugation (30000g, 1 h, 4 °C). The GST tag fused proteins were purified by affinity chromatography column with GSTrap HP (GE Healthcare) and the His or His-MBP tag fused proteins by nickel chelated HiTrap Chelating HP (GE Healthcare), followed by a TEV protease cleavage at 16°C overnight. A final step of size-exclusion chromatography was achieved using a Sephacryl S-100 HR 16/600 (GE Healthcare) with buffer Tris 50 mM (pH 7.5), NaCl 150 mM, TCEP 0.5 mM, protease inhibitors, one tablet per 100 mL of buffer (Complete, EDTA 2%, Roche Diagnostics). Regarding the purification of ZO-1-PDZ2, a denaturation protocol was applied. The bacterial pellet was resuspended in a denaturing solution containing guanidine hydrochloride (6M) and imidazole 20 mM, and cells were lysed by sonication. The debris were pelleted by centrifugation (30000g, 1 h, 4°C). The His-tagged ZO-1-PDZ2 were purified using a HiTrap Chelating HP (GE Healthcare), where a renaturation gradient by exchange of denaturation buffer to a gel filtration buffer (Tris 50 mM (pH 7.5), NaCl 150 mM, TCEP 0.5 mM, protease inhibitors, one tablet per 100 mL of buffer (Complete, EDTA 2%, Roche Diagnostics)) was performed. Protein elution was then carried out by a gradient of imidazole from 0 to 500 mM in gel filtration buffer. Finally, a size exclusion chromatography step was achieved using a Sephacryl® S-200 HR (GE Healthcare).

### 4.3 Peptide synthesis

The acetylated peptides containing the C-terminal PBM sequence of E protein (12 residues long) of SARS-CoV-2 (_ac_NLNSSRVPDLLV_COOH_) or of it South African mutant P71L (_ac_NLNSSRVLDLLV_COOH_) were synthesized in solid phase using the Fmoc strategy (Proteogenix, Schiltigheim, France). Peptides were resuspended in water to prepare stock solutions.

### 4.4 Crystallization, data collection, and structure determination

The PDZ – PBM complexes for crystallization were generated by mixing LNX2, MLLT4 or MPP5 PDZ domains with SARS-CoV-2 E protein PBM peptide at a ratio of 1:2, ensuring that at least 92% of complexes were formed. Initial screening of crystallization conditions was carried out by the vapor diffusion method using a MosquitoTM nanoliter-dispensing system (TTP Labtech) following the established protocols (Weber et al., 2019). Sitting drops were set up using 400 nL of a 1:1 mixture of each sample protein and crystallization solutions (672 different commercially available conditions) equilibrated against a 150 μL reservoir in multiwell plates (Greiner Bio-One). The crystallization plates were stored at 4 °C in a RockImager1000® (Formulatrix) automated imaging system to monitor crystal growth. The best crystals were obtained in crystallization conditions described in the table 2. Crystals were then flash cooled in liquid nitrogen using the condition of crystallization supplemented with 30% (V/V) of glycerol as cryoprotectant.

X-ray diffraction data were collected on the beamlines Proxima-1 and Proxima-2A at Synchrotron SOLEIL (St. Aubin, France). The data were processed by XDS (Kabsch, 2010), and the structures were solved by molecular replacement with PHASER (McCoy, 2007) using the search atomic models PDB id 3AXA, 4UU5 and 5E1Y for MLLT4-PDZ, MPP5-PDZ and LNX2-PDZ2, respectively. The positions of the bound peptides were determined from a Fo–Fc difference electron density maps. Models were rebuilt using COOT (Emsley et al., 2010), and refinement was done with phenix.refine of the PHENIX suite (Adams et al., 2010). The crystal parameters, data collection statistics, and final refinement statistics are shown in table 2. The structure factors and coordinates have been deposited in the Protein Data Bank under accession codes 7QCR, 7QCS, and 7QCT for MLLT4-PDZ, MPP5-PDZ, and LNX2-PDZ, respectively. All structural figures were generated with the PyMOL Molecular Graphics System, Version (Schrödinger).

### 4.5 GST Pulldown Assay

N-terminal GST fusion constructs containing the PDZ domains of ZO-1, MPP5, LNX2, PARD3 and MLLT4 were expressed and purified without the cleavage step by TEV protease. Purified GST constructs were individually incubated with glutathione-agarose beads for 1h at 4 °C with mild shaking. The beads were pelleted by centrifugation and washed 4 times with binding buffer (Tris 50 mM pH 7.5, NaCl 150 mM, TCEP 0.5 mM, antiprotease with EDTA 2% 1 tablet/100mL). HEK 293 cells were transiently transfected with the GFP tagged constructs using the phosphate calcium method. Cell lysates were prepared by scraping cells in lysis buffer Tris 50 mM pH7.5, triton 2%, NP40 1%, NaCl 200 mM with Complete protease inhibitor tablet (Roche, Indianapolis, IN) and centrifuged for 10 min at 13,000 rpm 4°C to pellet cell debris. Soluble detergent extracts were incubated with Glutathione resins for 2 hr at 4°C prior to washing three times with PBS supplemented with NaCl 200 mM and 0.1% Triton and processed for western blot analysis with GFP antibody (Novus NB600-313).

### 4.6 Immunofluorescence

HeLa cells were transfected using the Genejuice transfection reagent (Novagen) according to the manufacturer protocol. 24 hours after transfection, cells were fixed with PBS PFA 4% for 10 min and permeabilized in PBS Triton 0.1% for 5min before being processed for immunofluorescence using DAPI, anti-ALFA tag (NanoTag Biotechnologies; cat#N1502-SC3) and anti-GM130 (BD Transduction Lab; cat#610823). Images were acquired on a Leica DM6B microscope with a 63X objectives.

### 4.7 Microscale thermophoresis (MST)

The binding affinities between the C-terminal SARS-CoV-2-E PBM peptide (sequence NLNSSRVPDLLV_COOH_) or the P71L PBM peptide (sequence NLNSSRVLDLLV_COOH_) and the PDZ domains of ZO1, MLLT4, MPP5, LNX2 and PARD3 were measured using a Monolith NT.115 instrument (NanoTemper, Gmbh). The purified PDZ domains were covalently labelled using a fluorescent dye reactive on amine following the manufacture’s protocol (Protein Labelling Kit RED-NHS, Nanotemper). A serial dilution of the WT and the P71L mutant of the E protein PBM peptide was prepared in the buffer containing PBS tween 20 0.05%. A volume of 10 µl of peptide was serially diluted 1:1 in the buffer and mixed with an equal volume of labelled PDZ and loaded on capillaries. The concentration of the various labelled PDZ domains was kept constant and the peptide concentration varied.

We used 80% LED and 20% MST power at room temperature and MST time of 30 sec for all MST measurements. The data analysis and curve fitting with a Kd model were performed with NanoTemper programs MO.Control 2 and MO.Affinity 1 Analysis. The change in the thermophoretic mobility upon titration is measured as a delta of normalized fluorescence. All experiments were made in duplicate.

### 4.8 Analytical Ultracentrifugation (AUC)

ZO-1-PDZ2 and MLLT4-PDZ were prepared at 1 mg/ml. MLLT4-PDZ was prepared with an excess of protein E PBM peptide. Samples were prepared in Tris 50 mM pH 7.5, NaCl 150 mM, TCEP 0.5 mM. A sample of 400 µl were loaded into 1.2 cm double-sector cells between 2 sapphire windows. Cells were incubated for 2h at 20°C in an AN60-Ti rotor in the Optima-AUC analytical ultracentrifuge (Beckman Coulter) before data acquisition for 15h at 42,000 rpm. Sedimentation profiles were monitored over time by absorbance measurement at 280 nm and by interferometry. The data were analyzed using the continuous size distribution model c(s) of the Sedfit 16.36 software. All distributions were calculated with a floating frictional ratio f/f0 and a maximum entropy regularization procedure with a confidence level of 0.68. The buffer viscosity (η = 0.01031 Poise), the density (ρ = 1.0059) and the partial specific volume of 0.746 for MLLT4 and 0.736 for Z0-1 were estimated at 20°C from the amino acid sequences at 20 ° C using the software SEDTERP 3.0.3 (http://www.jphilo.mailway.com/sednterp.htm).

## 6 Author Contributions

CCS designed and directed the project. CCS and BB conceived and designed the experiments. YZ, FA, AH, SB, BN, BB, CCS collected the data. YZ, FA, AM, SB, BB, CCS analyzed the data. NW and SEM provided equipment and funding. CCS and BB wrote the manuscript with input from authors.

## 8 Acknowledgments

We thank the staff of the Crystallography core facility at the Institut Pasteur for carrying out robot-driven crystallization screenings, and the staff at the beamlines Proxima 1 and Proxima 2 at the French national synchrotron facility (SOLEIL, St Aubin, France), for help with data collection. We thank the DIM 1HEALTH region Ile-de-France for funding the Centrifection project that has allowed the Optima ultracentrifuge investment.

## 9 Supplementary Material

The Supplementary Material for this article can be found online at:

