## Supplementary figures and images for "Interactions of SARS-CoV-2 protein E with cell junctions and polarity PDZ-containing proteins"

### Supplementary Figure S1

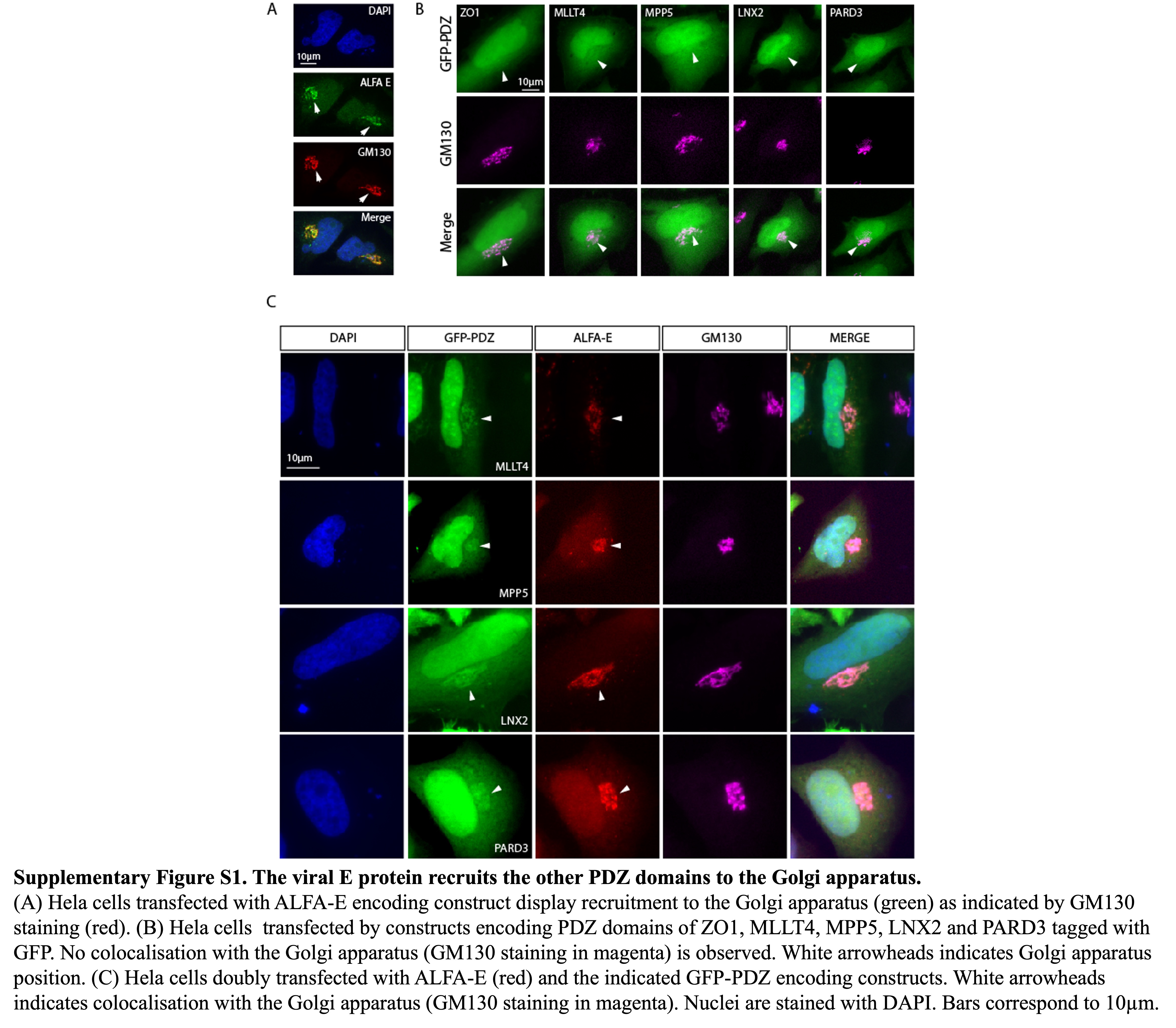

### Supplementary Figure S2

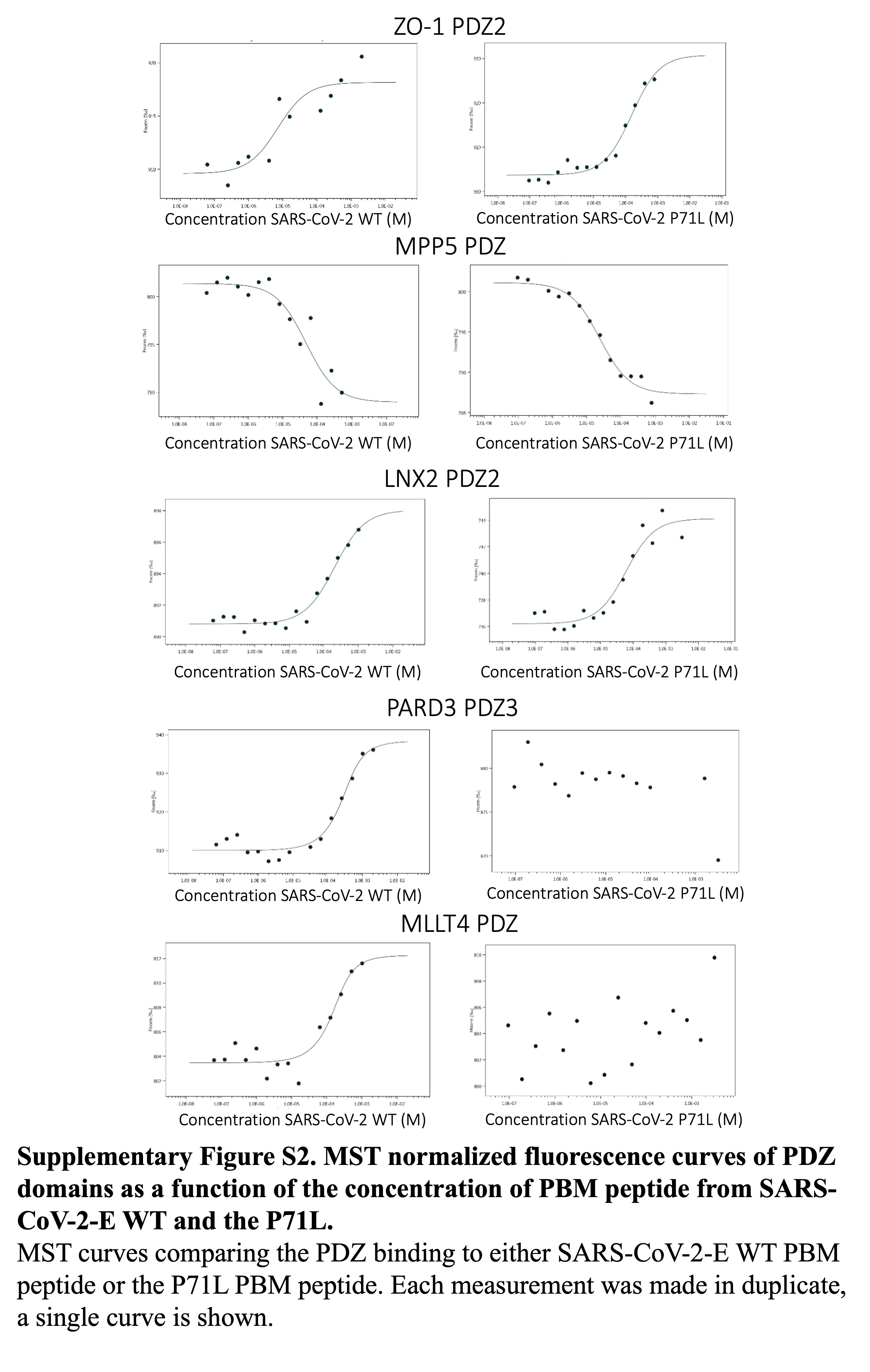

### Supplementary Figure S3

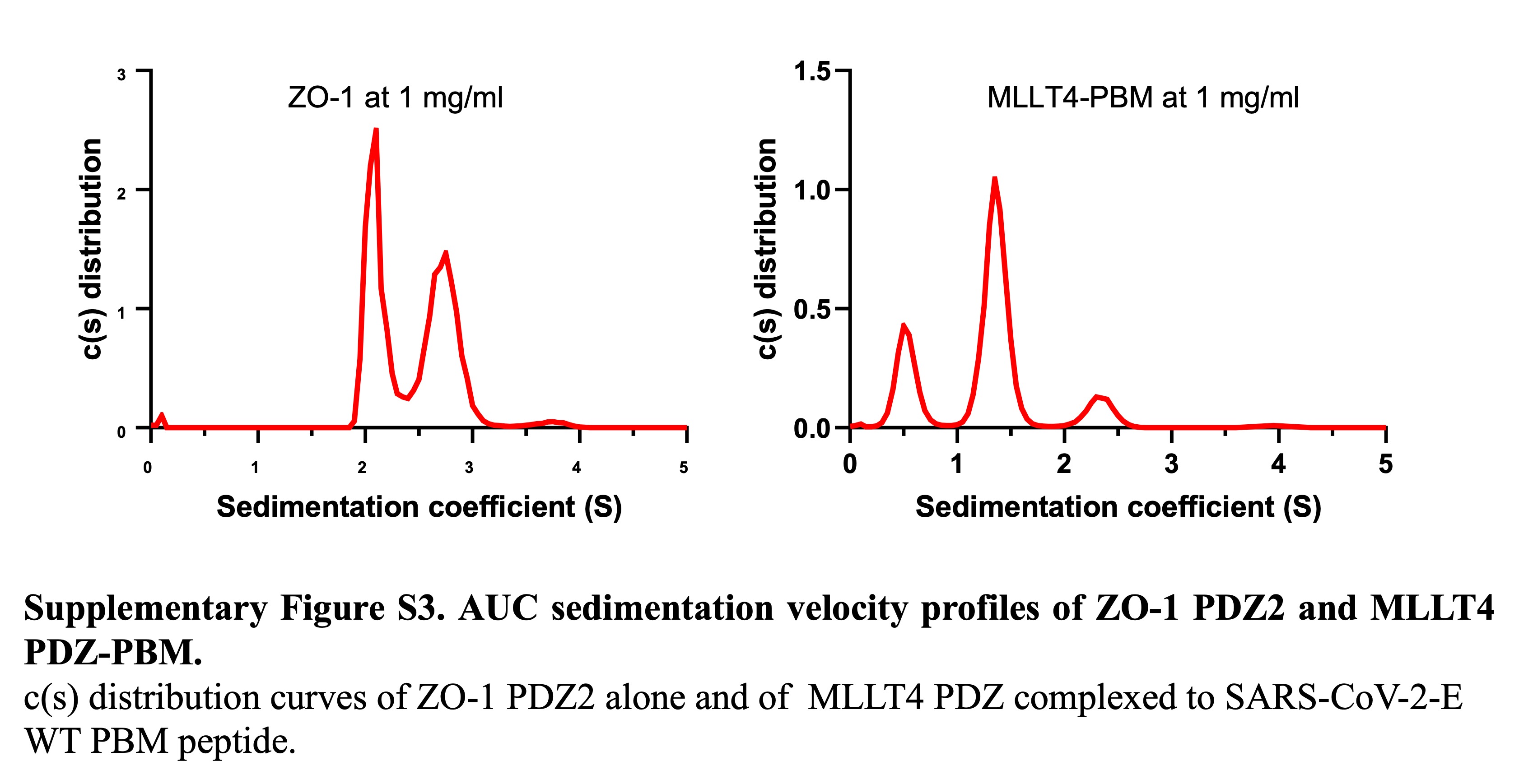
